# Biological Features Underlying Factor-Specific Predictability of Cross-Species Regulatory Occupancy

**DOI:** 10.64898/2026.04.14.718438

**Authors:** Yiman Wang, Guifen Liu, Yucheng Wang, Yong Zhang

## Abstract

Genome-wide maps of transcriptional and chromatin regulatory factor occupancy remain unevenly available across species, motivating computational approaches that leverage data from one species to predict regulatory occupancy in another. However, cross-species prediction performance varies substantially among regulatory factors, and the biological basis of this variability remains poorly understood. Here, we systematically evaluated cross-species predictability using 425 matched human–mouse ChIP-seq dataset pairs covering 137 regulatory factors. We identified diverse biological features associated with cross-species predictability, including motif architecture, evolutionary conservation, genomic context, and protein biophysical properties. These features collectively captured factor-specific differences in predictability and enabled accurate estimation of the probability that a regulatory-factor dataset is highly predictable across species. We further found that incorporating conservation-related information generally improved cross-species prediction, whereas additional regulatory context improved prediction for a subset of factors with weak or absent motif signals. Together, our study reveals a biological basis for factor-specific variation in cross-species regulatory occupancy prediction and provides a systematic framework for identifying factors amenable to cross-species prediction and determining when additional regulatory information is most informative.

## Introduction

Transcriptional and chromatin regulatory factors, including sequence-specific transcription factors (TFs), cofactors, and chromatin-associated proteins, play central roles in gene regulation and chromosome function [1–4]. Genome-wide mapping of their occupancy is therefore essential for elucidating regulatory mechanisms across diverse biological systems. Chromatin immunoprecipitation followed by sequencing (ChIP-seq) has enabled systematic characterization of regulatory-factor occupancy, but its application remains constrained by the availability and quality of suitable antibodies [5–9]. Epitope tagging provides an alternative for factors lacking suitable antibodies [10, 11], but engineering endogenous loci remains difficult to implement at scale. Consequently, regulatory-factor occupancy maps remain unevenly available across factors, tissues, and species.

Computational prediction provides an opportunity to extend regulatory annotations beyond experimentally profiled datasets. In particular, cross-species prediction leverages occupancy data available in one species to predict corresponding regulatory-factor occupancy in another, and recent studies have demonstrated the feasibility of this strategy using machine learning and deep learning approaches [12–17]. However, cross-species prediction performance varies substantially among regulatory factors. For example, CTCF can achieve AUPRC values of approximately 0.6, whereas GATA1 remains near 0.1 across different prediction methods [12, 13]. Such differences suggest that the feasibility of cross-species prediction depends strongly on the regulatory factor being modeled. Despite this pronounced variability, the biological properties associated with factor-specific cross-species predictability remain poorly understood.

Multiple aspects of regulatory-factor biology could contribute to these differences. For sequence-specific TFs, motif architecture and the extent to which in vivo binding sites contain recognizable motifs may influence how readily occupancy patterns can be learned from genomic sequence. Evolutionary conservation may also be important: CTCF, for example, has a highly conserved DNA-binding domain and shows conserved occupancy at many topologically associating domain boundaries across vertebrates [18–21]. Beyond intrinsic sequence specificity, regulatory occupancy is shaped by combinatorial interactions and chromatin environment. Cooperative TF interactions can alter sequence recognition and occupancy patterns [22, 23], while protein properties such as intrinsically disordered regions (IDRs) and phase-separation propensity may be associated with more context-dependent modes of chromatin association. Together, these observations suggest that cross-species predictability may reflect multiple aspects of regulatory-factor biology, ranging from sequence recognition and evolutionary conservation to regulatory context and protein biophysical properties.

Here, we systematically investigated factor-specific cross-species predictability using 425 matched human–mouse ChIP-seq dataset pairs covering 137 regulatory factors. We first quantified variation in cross-species prediction performance using a standardized sequence-based model and systematically identified biological features associated with this variability. We then developed a feature-based classifier to estimate the probability that individual regulatory-factor datasets are highly predictable across species. Finally, we examined whether incorporating conservation-related information and regulatory context could improve prediction and how the benefit of regulatory context varied among factors. Together, our study provides a systematic view of the biological features underlying factor-specific cross-species predictability and establishes a framework for determining when regulatory occupancy can be effectively predicted across species.

## Results

### Cross-species prediction performance varies systematically among regulatory factors

Previous studies have reported substantial variation in cross-species TF-binding site prediction performance [13] (Supplementary Figure S1A). To systematically characterize this variation across a broader range of regulatory factors, we assembled 425 human–mouse ChIP-seq dataset pairs manually matched by tissue or cell type, covering 137 transcriptional and chromatin regulatory factors (Supplementary Table S1). Each dataset contained more than 1,000 peaks. For each pair, we trained a sequence-based model, ChromTransfer-Base, in one species and evaluated it on the corresponding dataset in the other species, considering both mouse-to-human and human-to-mouse prediction (Figure 1A; see Methods for details). ChromTransfer-Base uses DNA sequence as input and applies the same model architecture and evaluation procedure across all regulatory-factor datasets, providing a standardized framework for comparing cross-species prediction performance.

**Figure 1.**
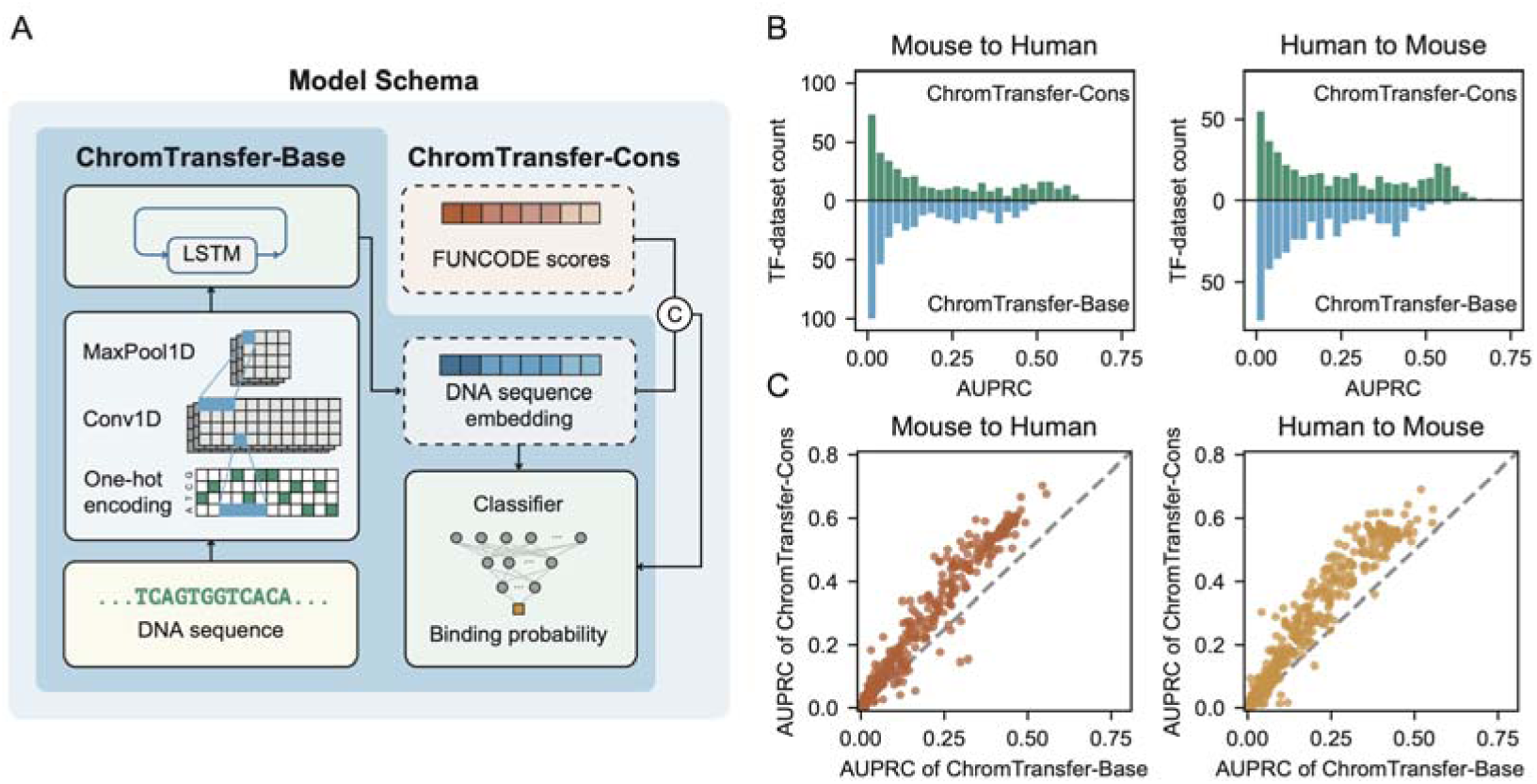
Factor-specific variation in cross-species regulatory occupancy prediction. (*A*) Schematic of the standardized cross-species prediction framework. ChromTransfer-Base predicts regulatory-factor occupancy from DNA sequence, whereas ChromTransfer-Cons additionally incorporates FUNCODE-derived regulatory correspondence scores. Models were trained in one species and evaluated on the matched dataset in the other species. (*B*) Distributions of AUPRC values for ChromTransfer-Base and ChromTransfer-Cons in mouse-to-human and human-to-mouse prediction. (*C*) Comparison of AUPRC values between ChromTransfer-Base and ChromTransfer-Cons in the two prediction directions. Mouse-to-human denotes training on mouse and evaluation on the matched human dataset; human-to-mouse denotes the reverse direction.

This systematic evaluation revealed substantial variation in cross-species predictability among regulatory factors. CTCF, RBBP5, and SUPT5H consistently showed high AUPRC values, whereas CBX3, NR3C1, and SNAI2 showed persistently low performance (Supplementary Figure S1B–D; Supplementary Table S2). The broad range of prediction performance observed under the same modeling and evaluation framework indicates that cross-species predictability differs markedly among regulatory factors. This variation motivated us to investigate the biological properties associated with factor-specific cross-species predictability.

We next examined whether incorporating additional information on regulatory conservation could improve cross-species prediction. To this end, we developed ChromTransfer-Cons, which extends ChromTransfer-Base by incorporating 12 FUNCODE-derived scores that capture cross-species regulatory correspondence [24] (Figure 1A; see Methods for details). ChromTransfer-Cons outperformed ChromTransfer-Base for most regulatory factors in both prediction directions (Figure 1B, C; Supplementary Figure S1B–D), indicating that regulatory correspondence information provides predictive value beyond DNA sequence alone. Nevertheless, substantial differences in prediction performance remained among regulatory factors, further supporting the need to understand the biological features associated with factor-specific cross-species predictability.

### Biological features associated with cross-species predictability

To investigate the biological basis of factor-specific variation in cross-species predictability, we analyzed 124 features in two broad categories (Supplementary Figure S2A; Supplementary Tables S3 and S4; see Methods for details). Peak-related features included motif properties, overlap with conserved and other genomic regions, repeat-associated measures, and co-occupancy with other regulatory factors, whereas protein-related features included DNA-binding-domain and whole-protein conservation, intrinsically disordered region (IDR) and sequence-composition measures, phase-separation-related properties, and protein interaction features. We calculated Pearson correlations between each feature and cross-species prediction performance, measured by AUPRC, across four combinations of model and prediction direction: ChromTransfer-Base and ChromTransfer-Cons in mouse-to-human and human-to-mouse prediction. Thirty-six features showed positive correlations above 0.2 across all four settings, whereas nine consistently showed negative correlations below −0.2 (Figure 2A). These results reveal multiple biological properties consistently associated with factor-specific cross-species predictability across model configurations and prediction directions.

**Figure 2.**
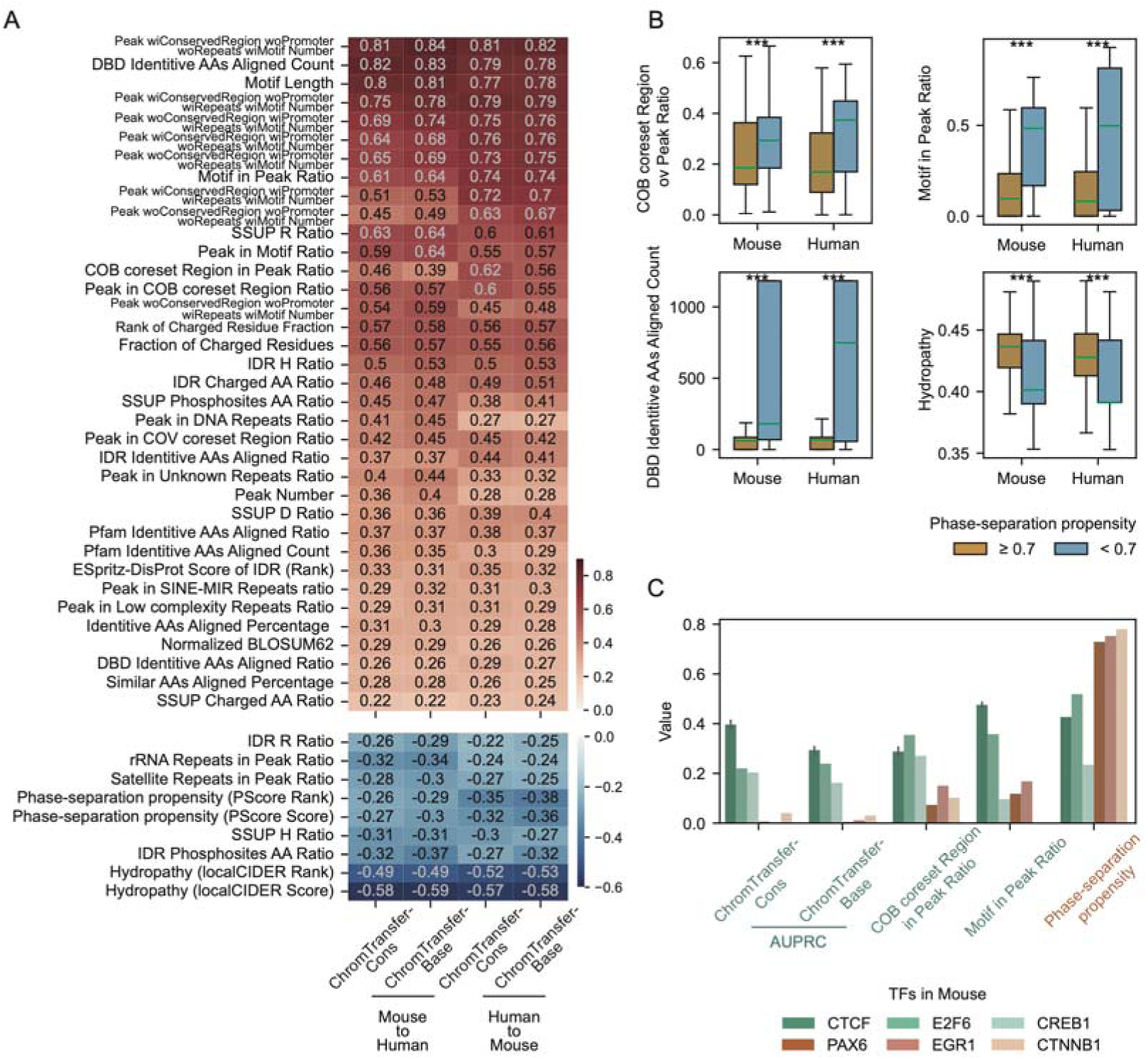
Biological features associated with cross-species regulatory-factor predictability. (A) Heatmap showing Pearson correlations between cross-species prediction performance and biological features across four combinations of model and prediction direction. Features with correlations greater than 0.2 (top) or less than −0.2 (bottom) across all four settings are shown. (B) Box plots of selected peak- and protein-related features for regulatory factors grouped by phase-separation propensity (PScore rank ≥0.7 or <0.7). Hydropathy represents the localCIDER score. (*C*) AUPRC values and selected biological features for representative regulatory-factor datasets in mouse. Green bars indicate factors with relatively higher prediction performance; reddish-orange bars indicate factors with relatively lower prediction performance.

Motif-related properties showed some of the strongest positive associations with cross-species prediction performance. In particular, motif length was among the most strongly correlated features across all four settings (Pearson r = 0.77–0.81), and factors whose ChIP-seq peaks more frequently overlapped their known motifs also tended to achieve higher AUPRC values (Figure 2A; Supplementary Figure S2B). Higher overlap with conserved regulatory regions and greater sequence conservation of DNA-binding domains between human and mouse orthologs were also positively associated with prediction performance (Figure 2A; Supplementary Figure S2B). Repeat-associated properties showed more heterogeneous relationships: overlap with specific SINE elements, particularly SINE-MIR, was positively associated with performance, whereas overlap with several other repeat classes, including rRNA and satellite repeats, showed negative associations. Together, these patterns indicate that regulatory factors characterized by stronger and more recognizable sequence determinants and greater evolutionary conservation tend to be more predictable across species.

Several protein and biophysical properties, by contrast, were negatively associated with cross-species prediction performance. Factors with higher phase-separation propensity, as measured by PScore [25], tended to have lower AUPRC values, with similar negative associations observed for IDR content, hydropathy, and several sequence-composition properties (Figure 2A; Supplementary Figure S2C). To further characterize this relationship, we divided regulatory factors according to PScore rank (≥0.7 or <0.7) [25, 26]. Factors with higher phase-separation propensity showed lower conserved-region overlap and motif content within their ChIP-seq peaks, fewer identical amino acids within DNA-binding domains between human and mouse orthologs, and higher hydropathy (Figure 2B). Representative factors showed a similar pattern: CTCF, E2F6, and CREB1 had lower PScore values together with higher prediction performance, motif content, and conserved-region overlap than PAX6, EGR1, and CTNNB1 (Figure 2C; Supplementary Figure S2D). These observations suggest that phase-separation-related properties are associated with a broader set of characteristics indicative of less sequence-constrained regulatory occupancy, which may contribute to reduced cross-species predictability.

Because many of the examined features are correlated with one another, these associations should not be interpreted as independent effects of individual features. Rather, they collectively indicate that cross-species predictability reflects multiple interconnected aspects of regulatory-factor biology, with motif architecture and evolutionary conservation associated with higher predictability and context-related protein properties associated with lower predictability.

### Biological features predict cross-species predictability

We next asked whether the biological features associated with cross-species prediction performance could be used to identify regulatory-factor datasets that are likely to be highly predictable. To this end, we developed an XGBoost-based classification framework using peak- and protein-related features as inputs (Figure 3A; see Methods for details). A dataset was defined as highly predictable if its cross-species prediction achieved both AUROC > 0.8 and normalized AUPRC above the corresponding Otsu-derived threshold in at least one of ChromTransfer-Base or ChromTransfer-Cons (Supplementary Figure S3A). We selected features showing Pearson correlations greater than 0.2 or less than −0.2 with AUPRC and used the union of these features, comprising 84 features in total, as inputs to the classifier (Supplementary Figure S3B; Supplementary Table S5). Using factor-grouped fivefold cross-validation, the classifier achieved an AUROC of 0.877 (Figure 3B), indicating that biological properties of regulatory factors and their occupancy profiles can effectively distinguish datasets with high and low cross-species predictability.

**Figure 3.**
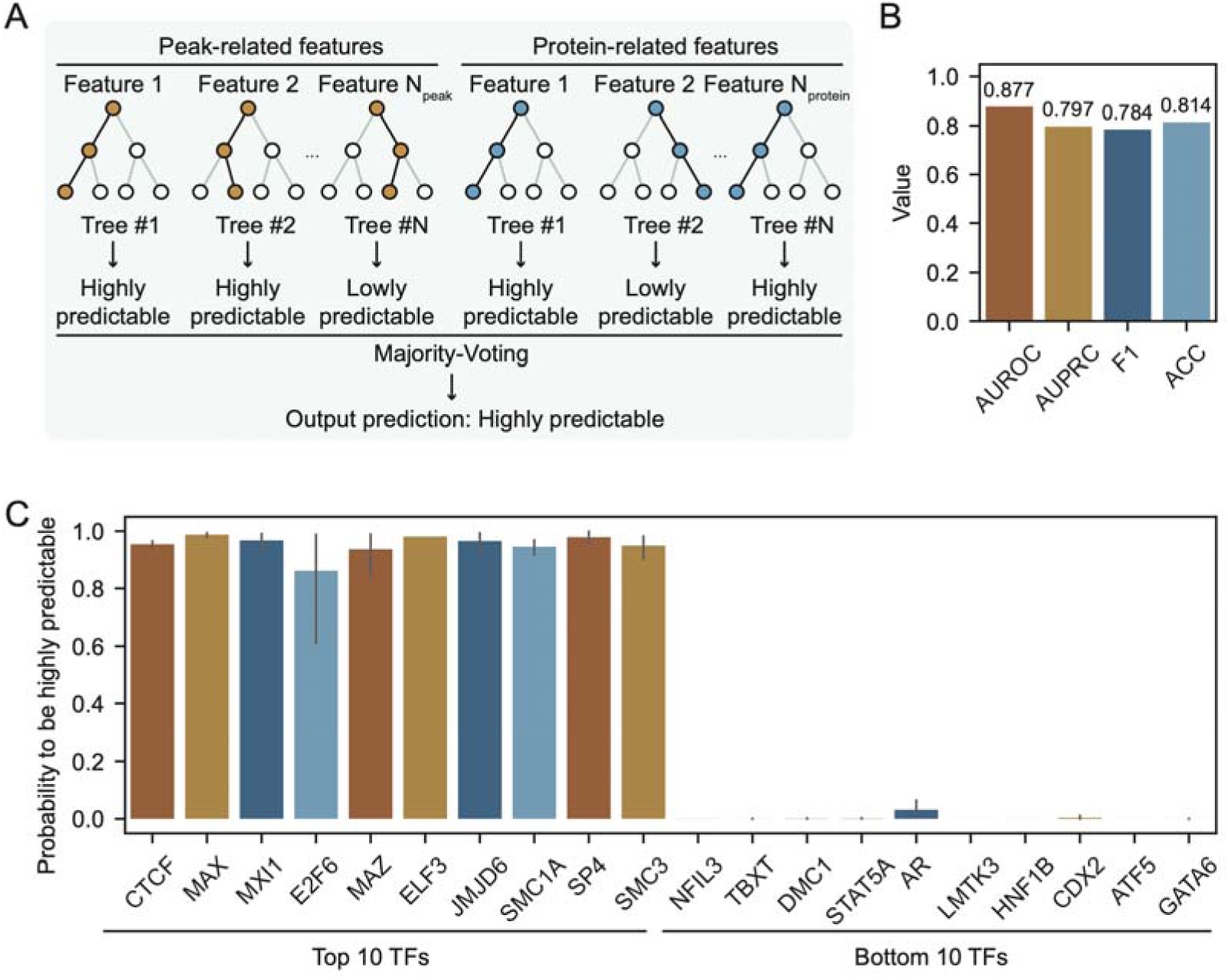
Feature-based classification of cross-species predictability. (*A*) Schematic of the XGBoost classification framework using peak- and protein-related features to classify regulatory-factor datasets as highly predictable or not highly predictable. The classifier assigns each dataset a probability of belonging to the highly predictable class. (*B*) Performance of the XGBoost classifier evaluated by factor-grouped fivefold cross-validation. (*C*) Predicted probabilities of being highly predictable for regulatory factors in the expanded dataset collection.

We next applied the classifier to an expanded collection comprising 2,728 datasets covering 723 regulatory factors in human and 1,915 datasets covering 238 regulatory factors in mouse (Supplementary Tables S6 and S7) [27]. The predicted probabilities of being highly predictable varied substantially among regulatory factors (Figure 3C; Supplementary Tables S6 and S7). CTCF, MXI1, SMC1A, and SMC3 received high probabilities, consistent with their strong performance in the matched-dataset analysis, while ELF3, JMJD6, and SP4, which were not included in classifier training, were also assigned high probabilities. In contrast, GATA6, AR, STAT5A, DMC1, and NFIL3 received low probabilities, as did ATF5, CDX2, HNF1B, and LMTK3, which were not included in classifier training. Interestingly, members of the GATA family, including GATA1, GATA2, GATA3, GATA4, and GATA6, shared similar feature profiles characterized by low proportions of motif-containing peaks and high phase-separation propensity, and were consistently assigned low probabilities of being highly predictable (Supplementary Figure S3C). By contrast, E2F6 and E2F2 differed substantially in their predicted probabilities despite belonging to the same family, accompanied by differences in motif-associated features (Supplementary Figure S3D). Together, these results show that biological features associated with cross-species prediction performance can also be used to identify regulatory-factor datasets that are more or less amenable to cross-species prediction.

### Regulatory context preferentially improves factors with weak sequence determinants

Motif-related properties, particularly motif length and the fraction of ChIP-seq peaks overlapping known motifs, were strongly associated with cross-species prediction performance (Figure 2A). Although regulatory factors with low cross-species predictability (AUPRC < 0.2) were as likely as highly predictable factors (AUPRC ≥ 0.2) to have at least one known motif, their peaks showed substantially lower overlap with these motifs (Supplementary Figure S4A). We therefore asked whether incorporating regulatory context could improve prediction for factors whose occupancy is less effectively captured by sequence information. To this end, we developed ChromTransfer-Reg, which extends ChromTransfer-Cons by incorporating two complementary categories of regulatory signals (Figure 4A; see Methods for details). Factor-specific regulatory signals were derived from occupancy profiles of candidate partner factors selected based on protein interaction evidence from STRING, co-occupancy information from ChIP-Atlas, or DNA-guided TF interaction evidence from CAP-SELEX experiments [22, 28, 29] (Supplementary Tables S1 and S8). Shared chromatin-context signals included matched chromatin accessibility profiles and 16 histone modifications (Supplementary Table S9). Datasets corresponding to the target factor itself were excluded from these regulatory inputs to avoid information leakage.

**Figure 4.**
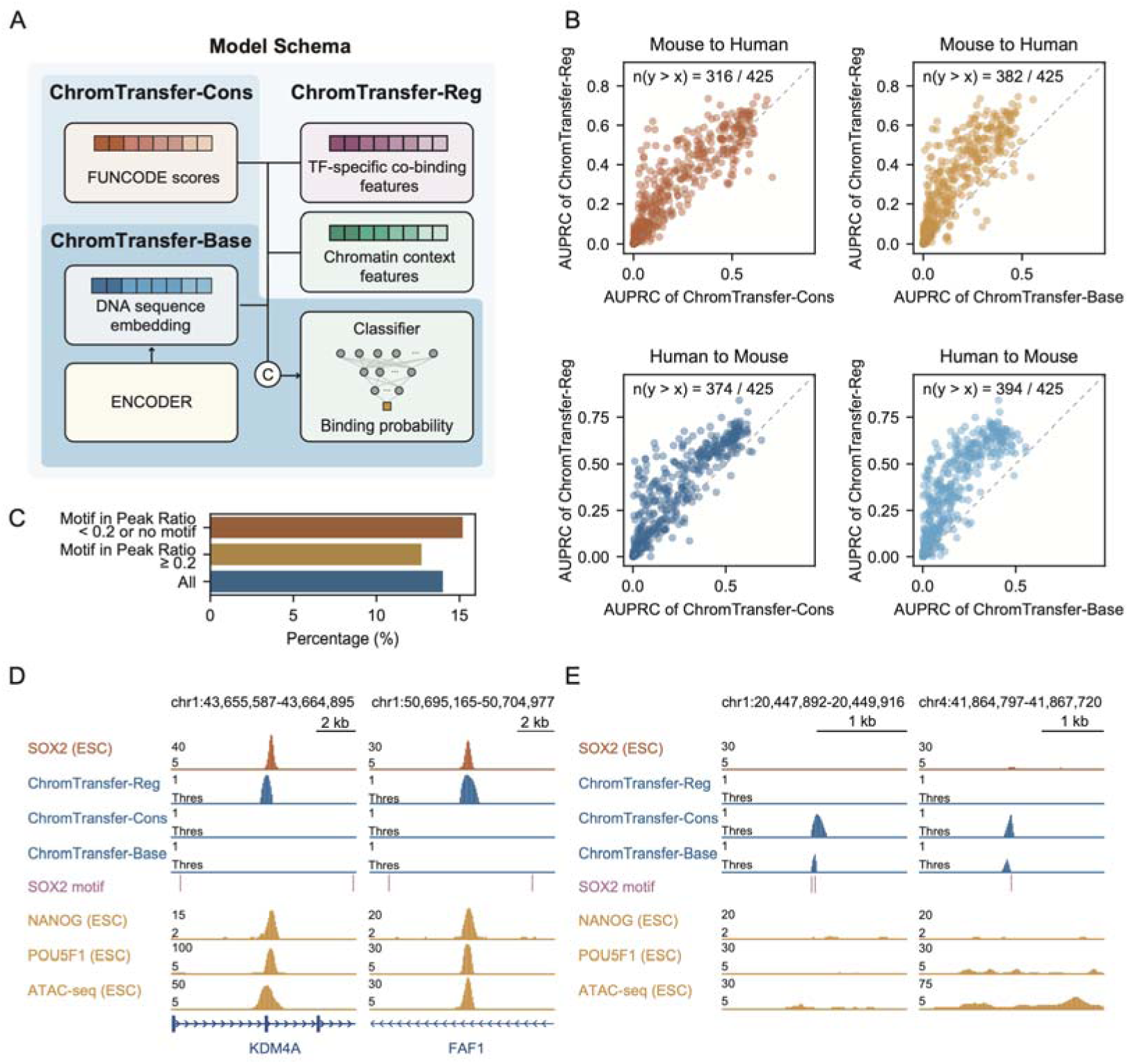
Regulatory context preferentially improves cross-species prediction for factors with weak sequence determinants. (*A*) Schematic of ChromTransfer-Reg, which extends ChromTransfer-Cons by incorporating factor-specific partner occupancy and shared chromatin-context signals. (*B*) Comparison of AUPRC values between ChromTransfer-Reg and ChromTransfer-Cons (left) or ChromTransfer-Base (right) in mouse-to-human and human-to-mouse prediction. (*C*) Percentage of datasets showing an AUPRC increase greater than 0.2 for ChromTransfer-Reg relative to ChromTransfer-Cons, stratified by motif–peak overlap (<0.2 or no known motif; ≥0.2; all datasets). (*D*) Representative human genomic regions containing SOX2 sites recovered by ChromTransfer-Reg, showing SOX2, NANOG, POU5F1, and ATAC-seq signals, model prediction scores, and SOX2 motif positions. (*E*) Representative false-positive sites predicted by ChromTransfer-Base and ChromTransfer-Cons but not by ChromTransfer-Reg.

ChromTransfer-Reg achieved higher AUPRC values than ChromTransfer-Cons and ChromTransfer-Base for most datasets in both prediction directions (Figure 4B), demonstrating that regulatory context provides additional predictive information beyond sequence and FUNCODE-derived regulatory correspondence. Importantly, however, the magnitude of improvement varied substantially among regulatory factors. Among datasets for factors lacking known motifs or with motif–peak overlap below 0.2, 15.2% showed an AUPRC increase greater than 0.2 relative to ChromTransfer-Cons (Figure 4C). Large performance gains were also more frequently observed for factors with access to more than 50 partner ChIP-seq dataset pairs (Supplementary Figure S4B). These results indicate that the contribution of regulatory context is factor-dependent, with particularly large gains observed for a subset of factors characterized by weak motif-associated sequence signals and rich partner occupancy information.

SOX2 provided a representative example of this pattern. For embryonic stem cells, ChromTransfer-Reg trained on mouse and evaluated on human improved AUPRC by 0.38 relative to ChromTransfer-Cons and by 0.41 relative to ChromTransfer-Base. The mouse SOX2 dataset had a low motif–peak overlap ratio of 0.143 and was accompanied by 58 partner ChIP-seq dataset pairs, including occupancy profiles for key regulators such as NANOG and POU5F1 (Supplementary Figure S4C). Sites missed by ChromTransfer-Base and ChromTransfer-Cons but recovered by ChromTransfer-Reg generally lacked SOX2 motifs while showing strong occupancy signals from partner factors (Figure 4D). Conversely, false-positive sites predicted by the sequence-based models frequently contained SOX2 motifs but lacked corresponding partner occupancy signals (Figure 4E). Thus, regulatory context can complement motif-based sequence information and substantially improve cross-species occupancy prediction for factors whose binding patterns are poorly captured by sequence alone.

## Discussion

Our study reveals substantial factor-specific variation in cross-species regulatory occupancy predictability and provides a systematic analysis of the biological properties associated with this variation. Using 425 matched human–mouse ChIP-seq dataset pairs covering 137 regulatory factors, we found that cross-species prediction performance varies markedly even when factors are evaluated using the same modeling framework. This variation was consistently associated with multiple aspects of regulatory-factor biology, including motif architecture, evolutionary conservation, genomic context, and protein biophysical properties. Moreover, these biological features could be used to identify regulatory-factor datasets with a high probability of successful cross-species prediction. Together, these findings suggest that cross-species predictability is not simply a property of the prediction model, but reflects systematic differences in how regulatory-factor occupancy is encoded by sequence, evolutionary constraints, and regulatory context.

Motif architecture emerged as one of the strongest features associated with cross-species predictability. Regulatory factors with longer motifs and greater motif–peak overlap generally showed higher prediction performance, consistent with the idea that distinctive sequence determinants provide stronger information for transferring occupancy patterns across species. CTCF represents a prominent example: its long and recognizable motif, high motif occupancy, conserved DNA-binding domain, and evolutionarily conserved genomic occupancy likely collectively contribute to its consistently high cross-species predictability. Beyond motif properties, higher overlap with conserved regulatory regions and greater conservation of DNA-binding domains were also associated with higher performance, further supporting a relationship between evolutionary constraint and cross-species predictability. By contrast, repeat-associated properties showed heterogeneous relationships with performance, suggesting that the contribution of repetitive elements depends on repeat class and regulatory context rather than following a uniform pattern.

Our results also point to a complementary relationship between sequence dependence and the biophysical properties of regulatory factors. Higher phase-separation propensity, IDR content, hydropathy, and several sequence-composition properties were associated with lower cross-species prediction performance. Recent studies have highlighted how IDRs and phase-separation-related interactions can modulate regulatory-factor occupancy and specificity beyond canonical DNA motif recognition [30, 31]. Consistent with this view, factors with higher phase-separation propensity in our analysis also tended to show lower motif–peak overlap and lower conservation-related measures. These correlated associations do not establish a direct causal effect of phase separation on cross-species predictability. Rather, they suggest that regulatory factors whose genomic occupancy depends more strongly on combinatorial interactions and local chromatin environments may be less readily captured and transferred by sequence-based models. This interpretation is also consistent with the improvement observed after incorporating partner occupancy and chromatin-context information, particularly for factors with weak motif-associated sequence signals.

The factor-specific nature of predictability has practical implications for cross-species regulatory modeling. Our feature-based classifier showed that biological properties of regulatory factors and their occupancy profiles contain sufficient information to distinguish datasets with relatively high and low cross-species predictability. Such estimates can provide a useful reference for prioritizing cross-species prediction tasks and for anticipating when sequence-based prediction is likely to be insufficient. The regulatory-context analysis further showed that the benefit of additional information is itself factor-dependent: regulatory context produced particularly large improvements for a subset of factors with weak motif-associated signals and extensive partner-occupancy information. Thus, rather than applying increasingly complex models uniformly to all regulatory factors, cross-species prediction strategies may benefit from considering the biological characteristics of the factor and the regulatory information available for the relevant cellular context.

Several limitations should be considered when interpreting these results. First, cross-species prediction performance reflects both the general predictability of regulatory-factor occupancy and the additional challenge of transferring predictions between species; the present analysis does not explicitly separate these two components. Future comparisons of within- and cross-species performance could help distinguish properties associated with general occupancy predictability from those specifically associated with cross-species transfer. Second, many of the biological features examined here are correlated, and their univariate associations therefore should not be interpreted as independent causal effects. Multivariate analyses across larger and more diverse datasets will be important for disentangling their relative contributions. Third, matching tissues and cell types between human and mouse is inherently imperfect, and differences in cellular composition or experimental context may contribute to the observed variation. Finally, incorporating regulatory context requires additional occupancy or chromatin datasets in the target species, which may limit its applicability in poorly characterized organisms. As regulatory datasets continue to expand across species and cellular contexts, integrating sequence, evolutionary, regulatory, and protein-level information should enable a more complete understanding of the factors governing cross-species regulatory occupancy.

## Methods

### Data collection and preprocessing

We collected paired human and mouse ChIP-seq datasets for transcriptional and chromatin regulatory factors from ChIP-Atlas, Cistrome Data Browser, and ENCODE [29, 32, 33]. The analyzed collection included sequence-specific transcription factors (TFs) as well as other transcriptional and chromatin-associated regulatory proteins; these proteins are collectively referred to as regulatory factors throughout the study. Human and mouse datasets were manually matched according to corresponding tissue or cell type. Only dataset pairs containing more than 1,000 peaks in both species were retained, resulting in 425 human–mouse dataset pairs covering 137 regulatory factors (Supplementary Table S1).

To standardize genomic inputs across species, the human and mouse genomes were divided into overlapping 500-bp bins with a stride of 50 bp. Bins overlapping ENCODE blacklist regions, which are prone to artifactual sequencing signals, were excluded using BEDTools intersect [33, 34]. For each regulatory-factor dataset, the remaining bins were assigned binary occupancy labels according to overlap with the corresponding ChIP-seq peaks and were used for model training and evaluation.

### Standardized cross-species occupancy prediction

To systematically quantify cross-species prediction performance, we used a sequence-based model, termed ChromTransfer-Base. The model takes DNA sequence as input and consists of a convolutional neural network (CNN) for detecting local sequence patterns, followed by a long short-term memory (LSTM) module and fully connected layers that output the probability that a genomic region is occupied by the target regulatory factor (Figure 1A).

For each matched human–mouse dataset pair, the model was trained using data from one species and evaluated on the corresponding dataset in the other species. Prediction was performed in both mouse-to-human and human-to-mouse directions. Throughout the study, cross-species predictability refers operationally to the prediction performance obtained when a model trained on one species is evaluated on the matched regulatory-factor dataset in the other species.

The area under the precision–recall curve (AUPRC) was used as the primary performance metric because regulatory-factor occupancy datasets contain substantially more unbound than bound genomic regions. AUROC was additionally calculated where indicated.

Chromosome 2 was held out as the test set. A random sample of 100,000 bins from chromosome 1 was used for validation, and all remaining eligible bins were used for model training. During each training epoch, all positive training bins were combined with an equal number of randomly sampled negative bins. Cross-species AUPRC was calculated using chromosome 2 of the target species. Dataset-level prediction results are provided in Supplementary Table S2.

### Incorporation of FUNCODE-derived regulatory correspondence information

To examine whether cross-species regulatory correspondence information provides predictive value beyond DNA sequence, we constructed ChromTransfer-Cons by extending ChromTransfer-Base with 12 FUNCODE-derived score tracks [24]. FUNCODE evaluates correspondence between human and mouse regulatory elements by integrating cross-species element relationships with functional genomic activity, including chromatin accessibility and histone modification profiles. We therefore refer to these inputs as FUNCODE-derived regulatory correspondence scores rather than measures of sequence conservation alone.

For each of the 12 FUNCODE BigWig tracks, the mean score within each 500-bp genomic bin was calculated, yielding a 12-dimensional feature vector. This vector was concatenated with the sequence representation generated by the sequence-processing layers and subsequently passed through multilayer perceptron (MLP) classification layers. ChromTransfer-Cons was trained and evaluated using the same cross-species procedure as ChromTransfer-Base.

### Calculation of biological features

We calculated 124 biological features representing characteristics of regulatory-factor occupancy profiles and properties of the corresponding proteins (Supplementary Tables S3 and S4). These features were broadly classified as peak-related features and protein-related features.

1. Peak number and motif-related features: The total number of ChIP-seq peaks in each dataset was obtained from the corresponding source database and included as a dataset-level feature. For sequence-specific TFs with available binding motifs, motifs were collected from CIS-BP [35], HOCOMOCO [36], JASPAR [37], UniPROBE [38], Wei *et al*. [39], Chen *et al*. [40], and TFBSshape [41]. When multiple motifs were available for a TF, motif length was averaged across motifs, weighted by the frequency with which each motif occurred within ChIP-seq peaks. Genome-wide motif occurrences were identified using FIMO from the MEME Suite [42]. Two complementary motif–occupancy features were then calculated using BEDTools intersect [34]: the fraction of ChIP-seq peaks containing the corresponding motif, termed Motif in Peak Ratio, and the fraction of motif occurrences overlapping ChIP-seq peaks, termed Peak in Motif Ratio.
2. Conservation- and FUNCODE-related features: Sequence conservation at regulatory-factor occupancy sites was quantified using PhastCons4way tracks for hg38 and mm10 obtained from the UCSC Genome Browser [43]. For each dataset, the mean PhastCons4way score across ChIP-seq peak regions was calculated and recorded as PhastCons4way. We also quantified overlap between ChIP-seq peaks and FUNCODE-defined region sets [24]. For the CO-B core set, CO-V core set, and CO-V extended set, we calculated both the fraction of peaks overlapping each region set and the fraction of each region set overlapping regulatory-factor peaks. These features retained their original labels, including COB coreset Region in Peak Ratio, COV coreset Region in Peak Ratio, COV extendset Region in Peak Ratio, and their reciprocal peak-in-region measures.
3. Repeat-associated features: Associations between regulatory-factor occupancy and repetitive elements were quantified using repeat annotations obtained from the UCSC Genome Browser [43]. For each repeat class, we calculated two complementary measures: the fraction of ChIP-seq peaks overlapping the corresponding repeat elements (Repeat in Peak Ratio) and the fraction of repeat elements overlapped by ChIP-seq peaks (Peak in Repeat Ratio). The repeat classes analyzed included low-complexity, scRNA, simple repeat, srpRNA, LINE, RNA, SINE, retroposon, tRNA, LTR, RC, satellite, rRNA, snRNA, DNA, and unknown repeats. We additionally examined the SINE subfamilies MIR, 5S-Deu-L2, tRNA-RTER, tRNA-Deu, Alu, and tRNA. For each subfamily, we calculated the fraction of peaks overlapping the subfamily and the fraction of subfamily elements overlapped by peaks.
4. Composite genomic-context features: To characterize combinations of genomic contexts, we constructed 16 composite features representing all possible combinations of presence or absence of overlap with four categories of genomic elements: FUNCODE-defined regions, promoters, repeat elements, and motifs. For each combination, we calculated the proportion of ChIP-seq peaks satisfying the corresponding overlap pattern. For example, peak wiConservedRegion wiPromoter wiRepeats wiMotif Ratio represents the fraction of peaks overlapping all four categories.
5. Regulatory-factor co-occupancy: Genomic co-occupancy of each regulatory factor with other factors was calculated using human and mouse co-occupancy matrices for 1-kb genomic regions derived from the ChromBERT dataset [27]. For each target regulatory factor, we counted co-occupancy events with other factors and averaged these counts across available datasets after excluding datasets corresponding to the target factor itself. Because these measurements are based on overlap of genomic occupancy profiles, they are referred to as co-occupancy rather than direct physical co-binding.
6. Protein-association features: Protein-association information was obtained from STRING [28]. Sequence-specific TFs were identified using AnimalTFDB 4.0 [44], and chromatin regulators were identified using CRdb [45]. For each applicable regulatory factor, we calculated the number of STRING associations with a combined score >700 and the mean STRING combined score across its retained associations. These features retained their original labels, Number of TF–TF interactions and Average TF–TF interaction score.
7. Phase-separation-related features: Phase-separation propensity was characterized using multiple computational approaches. PSPire was used to obtain an overall phase-separation propensity score and a score calculated while ignoring IDRs [46]. MolPhase scores were obtained from MolPhase [47]. Additional scores were obtained from PhaSePred [26], including catGRANULE score and rank, PScore score and rank, and SaPS-8fea and PdPS-8fea scores and ranks [25, 26, 48]. These features represent computational estimates of phase-separation-related properties rather than direct experimental measurements of condensate formation.
8. IDR, SSUP, and amino-acid-composition features: Intrinsically disordered regions (IDRs) were identified using IUPred2A with a score threshold of 0.5 and a minimum length of 20 amino acids [49]. Relative solvent-accessible surface area was calculated using PSAIA [50], and structured superficial regions (SSUPs) were defined following the procedure used in PSPire [46]. The resulting features included total IDR length, IDR fraction, fractions of charged amino acids and phosphorylation sites within IDRs and SSUPs, and the fractions of lysine, arginine, histidine, aspartic acid, and glutamic acid within these regions.
9. Protein sequence and domain conservation: Human–mouse protein sequence conservation was evaluated using BLOSUM62 alignment scores [51]. We calculated the raw alignment score, length-normalized alignment score, percentage of identical aligned residues, and percentage of similar residues, with similarity defined as an aligned residue pair with BLOSUM62 score >0. Conservation was additionally evaluated within Pfam-annotated domains [52], DNA-binding domains defined by Lambourne *et al*. [53] and IDRs. For each region type, both the number and fraction of identical aligned residues were calculated. Protein domains were identified using PyHMMER [54].
10. Additional protein biophysical features: Additional protein-sequence-derived features were obtained from PhaSePred [26], including PLAAC scores for prion-like domains [55], ESpritz-DisProt scores for intrinsic disorder [56], localCIDER hydropathy scores [57], DeepCoil coiled-coil scores [58], SEG low-complexity scores [59], and charged-residue fractions. Raw and rank-normalized versions were included where available. For regulatory factors with the required orthologous coding sequences, Ka/Ks ratios were calculated using KaKs_Calculator 3.0 [60]. Protein structural similarity between human and mouse orthologs was quantified using TM-align [61]. Protein structures were obtained from the AlphaFold Protein Structure Database [62]. The resulting TM-align score, ranging from 0 to 1, was included as a feature representing cross-species structural similarity.

### Association analysis of biological features with cross-species predictability

For each feature, Pearson correlation coefficients were calculated independently against cross-species prediction performance measured by AUPRC. Correlations were evaluated for four combinations of prediction model and direction: ChromTransfer-Base mouse-to-human, ChromTransfer-Base human-to-mouse, ChromTransfer-Cons mouse-to-human, and ChromTransfer-Cons human-to-mouse.

Features showing Pearson correlation coefficients >0.2 or <−0.2 were considered strongly associated with prediction performance for the analyses shown in Figure 2 and for subsequent feature selection. For the comparison shown in Figure 2B, regulatory factors were divided according to PScore rank into groups with rank ≥0.7 and <0.7.

### Feature-based classification of highly predictable datasets

To determine whether biological features could identify regulatory-factor datasets likely to perform well in cross-species prediction, we trained XGBoost classifiers [63]. For each combination of prediction model and direction, a dataset was labeled highly predictable if its cross-species prediction performance satisfied both AUROC >0.8 and normalized AUPRC above the corresponding Otsu-derived threshold (Otsu 1979).

AUPRC was normalized according to the proportion of positive samples:

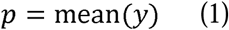

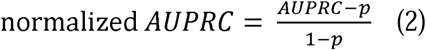

where p represents the positive-class proportion. Otsu thresholding was then applied to the normalized AUPRC distribution to separate datasets into higher- and lower-predictability groups.

Features with Pearson correlations >0.2 or <−0.2 with the corresponding cross-species AUPRC were retained, and the union of the selected features was used as classifier input, resulting in 84 features.

Four XGBoost classifiers were trained separately for ChromTransfer-Base mouse-to-human, ChromTransfer-Base human-to-mouse, ChromTransfer-Cons mouse-to-human, and ChromTransfer-Cons human-to-mouse prediction. Dataset splitting was grouped by regulatory factor such that datasets corresponding to the same factor were assigned to the same fold. Model performance was evaluated using fivefold GroupKFold cross-validation [64] (Supplementary Table S5). Hyperparameters were selected by cross-validated grid search, after which the classifiers were retrained using all available matched datasets for application to the expanded datasets. Each classifier produced the probability that an individual regulatory-factor ChIP-seq dataset belonged to the highly predictable class.

### Application of the classifier to expanded regulatory-factor datasets

Expanded collections of human and mouse ChIP-seq datasets were obtained from Yu *et al*. [27] (Supplementary Tables S6 and S7). After retaining datasets containing more than 1,000 peaks, the collection comprised 2,728 datasets covering 723 regulatory factors in human and 1,915 datasets covering 238 regulatory factors in mouse. The biological features required by the XGBoost classifiers were calculated for each dataset, and the trained classifiers were used to assign probabilities of belonging to the highly predictable class.

### Integration of partner occupancy and chromatin context

To examine whether additional regulatory information improves cross-species prediction for factors with weak sequence-associated signals, we constructed ChromTransfer-Reg by extending ChromTransfer-Cons with two types of regulatory inputs: factor-specific partner occupancy signals and shared chromatin-context signals.

Candidate partner factors were selected using three complementary sources of evidence. STRING protein associations with combined scores ≥700 were used to identify high-confidence protein associations [28]. ChIP-Atlas was used to identify factors showing strong genomic co-occupancy with the target regulatory factor, using an average score >4.5 [29]. CAP-SELEX provided experimental evidence for DNA-guided TF interactions [22]. For each selected partner, its ChIP-seq occupancy profile was used as the actual model input. Supplementary Table S8 lists the partner datasets used for each target regulatory factor.

Shared chromatin-context inputs consisted of matched human–mouse chromatin-accessibility datasets and 16 histone-modification datasets (Supplementary Table S9). Importantly, occupancy datasets corresponding to the target regulatory factor itself, irrespective of cell type, were excluded from the regulatory inputs.

BigWig and peak files were obtained from Cistrome Data Browser or ChIP-Atlas [29, 32]. Regulatory signals were discretized within 500-bp bins following the procedure used in ChromBERT [27]. The mean BigWig signal was first calculated for each bin. A bin was classified as peak-overlapping if more than 50% of a peak overlapped the bin or more than 80% of the bin overlapped a peak. Non-peak bins with mean signal below or above the median of the corresponding BigWig track were assigned values of 0 or 1, respectively. Peak-overlapping bins were divided into tertiles according to their mean signal and assigned values of 2, 3, or 4 from lowest to highest tertile.

The resulting partner-occupancy and chromatin-context signals were combined into a regulatory-context vector and concatenated with the sequence representation and FUNCODE-derived scores before the MLP classification layers (Figure 4A).

### Genome browser visualization

Genome-browser views were generated using the UCSC Genome Browser and track data hubs [43, 65]. Prediction probabilities from overlapping 500-bp bins generated at 50-bp intervals were averaged to produce 50-bp-resolution signal tracks. Only signals above the selected prediction thresholds were displayed. Prediction thresholds were selected to maximize F1 scores from precision–recall curves in the target species.

### Statistical analysis

Unless otherwise specified in the figure legends, comparisons between groups were performed using two-sided Mann–Whitney U tests implemented in SciPy [66]. Statistical significance is denoted as ***P<0.001; n.s., not significant.

### Code availability

Regulatory-factor co-occupancy and chromatin-context signals for 500-bp genomic regions with a 50-bp step size in mouse (mm10) and human (hg38) are available on Figshare at: https://doi.org/10.6084/m9.figshare.31972113. Source codes of the cross-species prediction models are available on the GitHub repository (https://github.com/TongjiZhanglab/ChromTransfer) and Zenodo repository (https://zenodo.org/records/19449463).

## Acknowledgements

This work was supported by the National Natural Science Foundation of China (32488101, 32325012), the National Key Research and Development Program of China (2021YFA1302500), the Science and Technology Commission of Shanghai Municipality (23JS1401200, 26XF3200400).

## Author contributions

Y.Z. conceived the project; Y(Yiman).W., G.L. and Y(Yucheng).W. conducted data collection, model development, and feature analysis; Y.Z., Y(Yiman).W., and G.L. wrote the manuscript.

